# Athlytics: A Computational Framework for Longitudinal Analysis of Exercise Physiology Metrics from Wearable Sensor Data

**DOI:** 10.1101/2025.05.01.651597

**Authors:** Zhiang He

## Abstract

The proliferation of wearable sensors provides large-scale longitudinal physiological data collected in real-world settings, offering unprecedented opportunities to investigate dynamic human responses to exercise interventions. However, systematically quantifying key physiological indicators related to adaptation and fatigue from these dense time-series data, particularly from popular platform APIs like Strava, and performing standardized, reproducible integrative analyses currently lacks established open-source workflows in R, posing significant practical and computational barriers. Researchers often expend considerable effort on custom programming, limiting analytical scale and efficiency. To overcome this critical bottleneck and empower broader research applications, we developed and introduce `Athlytics`, a computational framework specifically designed for processing Strava API data directly within R for longitudinal exercise physiology analysis. This framework provides a dedicated means to seamlessly integrate data acquisition with the calculation of key physiological indicators (such as ACWR reflecting training stress balance, EF assessing aerobic efficiency, and physiological decoupling indicating cardiovascular stability), based on activity summary data while carefully addressing necessary approximations for composite load metrics like TSS/HRSS, and multidimensional time-series visualization. By providing standardized function interfaces, `Athlytics` significantly lowers the technical barrier for conducting complex longitudinal analyses, enabling researchers to efficiently test hypotheses regarding the dynamic interplay between training stimuli, physiological efficiency, and stress responses. This work provides an open-source computational tool that fills a critical gap, contributing by substantially enhancing the feasibility, efficiency, and reproducibility of quantitative exercise physiology research utilizing widely available physiological sensor data through standardization and automation. The package (https://github.com/HzaCode/Athlytics) provides an important foundation for standardizing and applying computational methods in this field.

## 1 Introduction

Understanding the dynamic physiological responses to exercise stimuli is central to exercise science and personalized health [1]. The widespread adoption of wearable sensors and online platforms like Strava is generating vast streams of longitudinal physiological and behavioral data collected in real-world settings [2]. This data deluge presents significant opportunities for quantifying training loads, monitoring physiological adaptations, and potentially mitigating injury risks. However, translating this raw data into actionable, scientifically rigorous insights faces considerable methodological and computational challenges. Effective analysis requires not only accessing the data but also integrating diverse metrics over time, including those reflecting training stress balance (e.g., Acute:Chronic Workload Ratio, ACWR [3–5]), aerobic system efficiency (e.g., Efficiency Factor, EF [6]), performance milestones (e.g., personal bests, PBs [7]), and indicators of cardiovascular stability or fatigue (e.g., heart rate/power decoupling [8]).

While the theoretical basis for these metrics is established, their systematic, reproducible implementation within a standard computational environment like R [9], particularly using data directly sourced from heterogeneous platform APIs (such as Strava’s), remains problematic. Existing tools may lack specific functions for these exercise physiology indicators, or require substantial custom scripting to link data retrieval, calculation, and longitudinal visualization. This lack of standardized, open-source workflows constitutes a significant practical barrier, hindering the efficiency, scale, and reproducibility of research utilizing these ubiquitous data sources [10]. Researchers often dedicate considerable time to developing bespoke code, diverting effort from core scientific questions. Furthermore, reliance on readily available summary statistics from APIs necessitates careful evaluation of the validity of derived metrics, as approximations (e.g., estimating Training Stress Score (TSS) from average power rather than normalized power) can significantly impact physiological quantification, particularly when comparing sessions with varying intensity profiles [6]. Similarly, the inherent variability and potential inaccuracies in sensor-derived data (e.g., GPS-based pace/distance, user-provided thresholds) require cautious interpretation of results.

To address the practical barrier of accessing and analyzing these data streams within R, we developed `Athlytics`. This work introduces `Athlytics` not merely as a collection of functions, but as a **computational framework** designed to facilitate quantitative, reproducible research, while explicitly acknowledging the data source and methodological limitations. It specifically targets the workflow from Strava API data retrieval (via the `rStrava` package [11]) to the longitudinal analysis and visualization of key exercise physiology metrics. The framework operationalizes the calculation of ACWR, Load Exposure (visualizing acute vs. chronic load states), PB tracking, EF trends, and Decoupling patterns through dedicated modules. By adopting `tidyverse` principles [12] and providing paired `calculate_*` and `plot_*` functions adhering to `tidyverse` principles [12], `Athlytics` facilitates a structured workflow from raw data retrieval (via `rStrava`) to interpretable analytical outputs. This work aims to lower the barrier to conducting multi-faceted longitudinal training analysis using ubiquitous Strava data, thereby contributing a necessary tool to the sports science computational ecosystem, intended for use with a critical understanding of its assumptions.

This paper details the design of the `Athlytics` framework, its dependencies, and its core analytical modules. We elaborate on the methodological implementation within each module, including necessary assumptions. Through illustrative examples, we demonstrate its application to typical analytical questions in training monitoring. Finally, we position `Athlytics` within the landscape of related computational tools, dedicate significant discussion to the critical limitations imposed by data source quality and metric approximations, and outline future directions.

## 2 Overview of the Athlytics Package

The `Athlytics` package is designed as an R framework offering a concise, efficient, and userfriendly environment for analyzing personal sports training data sourced from Strava. It focuses on operationalizing the calculation and visualization of key performance and physiological load metrics, notably the Acute:Chronic Workload Ratio (ACWR), EF, and decoupling. The core philosophy centers on integrating data acquisition, standardized metric calculation, and results visualization, thereby minimizing the manual operations and bespoke programming typically involved. Adhering to `tidyverse` [12] design principles, `Athlytics` utilizes the pipe operator (`%>%`) via `dplyr` [13] to promote readable code and coherent analytical workflows.

### 2.1 Dependencies and Installation

`Athlytics` builds on a robust ecosystem of well-established R packages. For data acquisition, it utilizes `rStrava` [11] to handle API authentication and data retrieval. Core data wrangling is performed using `dplyr` [13] and `tidyr` [14], while date-time manipulation is supported by `lubridate` [15]. Rolling-window calculations for training load metrics are implemented via `zoo` [16]. Data visualization is powered by `ggplot2` [17]. In addition, `rlang` [18] and `purrr` [19] provide functional programming tools that enhance internal package logic.

`Athlytics` is available on CRAN and can be installed directly as follows:

**Listing 1:**
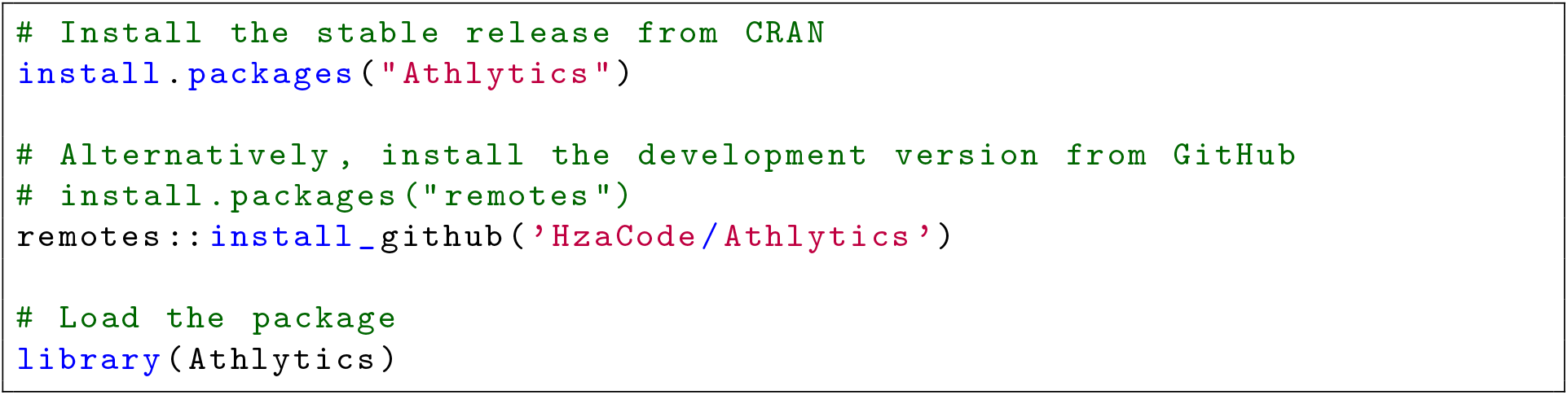
Installation options for the Athlytics package.

The CRAN version is recommended for most users. Advanced users seeking the latest development updates may opt for the GitHub version.

#### Core Analysis Modules

The framework provides distinct modules for key analytical tasks, each generally featuring a `calculate_*` function for computation and a `plot_*` function for visualization. This modular structure allows researchers flexibility in executing specific analyses or retrieving intermediate data outputs for custom downstream processing.

### 2.2 ACWR Trend

The Acute:Chronic Workload Ratio (ACWR) is a widely adopted method for monitoring training stress balance and identifying periods of potentially heightened injury risk based on rapid load changes. `Athlytics` implements the common rolling average approach [3].

#### Calculation Function

`calculate_acwr` Fetches activities via Strava API, filtered by activity type if specified. Calculates load based on the user-specified metric (duration, distance, TSS, etc.). It should be noted that when TSS or HRSS are selected, these are approximated based on summary statistics (e.g., average power/HR) rather than detailed time-series methodologies, a limitation discussed further in Section 4.2. Groups loads by day to create a continuous daily load time series. Computes acute (7-day) and chronic (28-day) rolling averages. Divides acute by chronic load to generate the ACWR time series.

#### Visualization Function

`plot_acwr` Creates a time series visualization of the ACWR. Visualizes the smoothed ACWR over time. Optionally shades background regions corresponding to commonly cited ACWR risk zones, adding reference lines. Generates informative titles and subtitles reflecting the chosen parameters. Outputs a standard `ggplot` object, allowing for further user customization.

### 2.3 Load Exposure

Visualizing the interplay between acute load (ATL) and chronic load (CTL) offers a complementary perspective on the athlete’s current state within the training load landscape [3]. The Load Exposure Plot facilitates rapid assessment relative to ACWR-derived thresholds.

#### Calculation Function calculate_exposure

Computes the necessary data for the exposure plot, following steps analogous to `calculate_acwr` but returning daily `atl`, `ctl`, and `acwr` values. When `load_metric = “tss”` is selected, the user must provide their Functional Threshold Power via the `user_ftp` argument; the scientific validity implications of using TSS approximated from summary statistics are discussed in Section 4.2.

#### Visualization Function plot_exposure

Generates the scatter plot using `calculate_exposure` output and `ggplot2` [17]. Plots ATL versus CTL, with each point denoting a day. Optionally incorporates diagonal rays or shaded areas indicating ACWR thresholds (e.g., 0.8, 1.3, 1.5) to delineate risk zones. The position of the most recent data point concerning these zones aids in evaluating the current load management status.

### 2.4 Personal Bests (PBs)

Tracking personal bests (PBs) provides milestones for performance progression, reflecting peak physiological output capabilities over specific durations or distances [7].

#### Calculation Function calculate_pbs

Scans Strava activities to identify and record best performance times for user-specified distances. Retrieves activities via `stoken`, filtering by `activity_type` and optional `date_range`. For each distance in `distance_meters`, analyzes activity data to determine the fastest time, relying on Strava’s potentially incomplete or inaccurate `best_efforts` data field available in the activity summary. The reliability of this source for rigorous research is discussed in Section 4.2. Returns a `tibble` summarizing activities where PBs were identified.

#### Visualization Function plot_pbs

Uses the output from `calculate_pbs` to visualize PB progression over time with `ggplot2` [17]. Plots completion time against date for each specified distance. Visually distinguishes points representing identified PBs. Facilitates comparison by displaying trends for multiple distances simultaneously.

### 2.5 Efficiency Factor (EF)

The Efficiency Factor (EF) serves as an indicator of aerobic efficiency, relating mechanical output (pace or power) to physiological cost (heart rate). Longitudinal tracking can reveal adaptations in the aerobic system or potential effects of fatigue [6].

#### Calculation Function calculate_ef

Computes EF metrics from Strava activities. Retrieves activities, with optional filtering based on minimum duration (`min_duration_mins`). Calculates EF based on the chosen `ef_metric` (`“Pace_HR”` : average speed / average heart rate; `“Power_HR”` : average power / average heart rate), requiring relevant data (average speed/pace, power, heart rate) within the activity summary. Returns a `tibble` containing `date`, `activity_type`, and the calculated `ef_value`.

#### Visualization Function plot_ef

Visualizes changes in EF over time using `ggplot2` [17]. Generates a scatter plot of EF against date. Optionally overlays a smoothing trend line to discern long-term patterns.

### 2.6 Decoupling

Heart rate/power (or pace/heart rate) decoupling quantifies the dissociation between physiological input (heart rate) and mechanical output (power or pace) during sustained aerobic exercise. Significant decoupling can indicate limitations in cardiovascular stability, substrate availability, or hydration status under prolonged stress [8, 20].

#### Calculation Function calculate_decoupling

Computes the decoupling rate for activities. Retrieves detailed activity streams (time-series data for heart rate, power, or speed) via `stoken`. This approach requires appropriate API permissions (`activity:read`), significantly increases processing time, and consumes more API quota compared to fetching summary data. Segments the activity data into first and second halves (typically by duration). Determines the efficiency ratio (output/HR) for each half. Calculates the percentage difference: `(First Half Ratio - Second Half Ratio) / First Half Ratio * 100%`. Returns a data structure linking activity identifiers with the calculated `decoupling_percent`.

#### Visualization Function plot_decoupling

Visualizes decoupling trends. Plots the calculated `decoupling_percent` over time across multiple activities using `ggplot2` [17].

## 3 Illustrative Applications

This section illustrates the application of the `Athlytics` framework to address typical analytical questions using its core modules. The figures presented below were generated using the package’s plotting functions (with sample data) and demonstrate the type of insights that can be efficiently generated.

**Listing 2:**
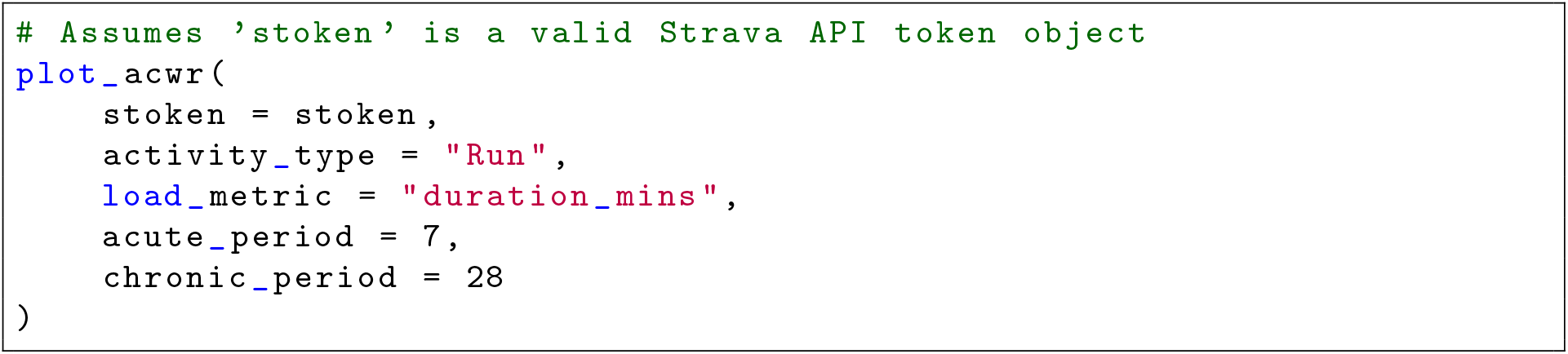
R code generating the ACWR trend plot (Figure 1).

**Figure 1:**
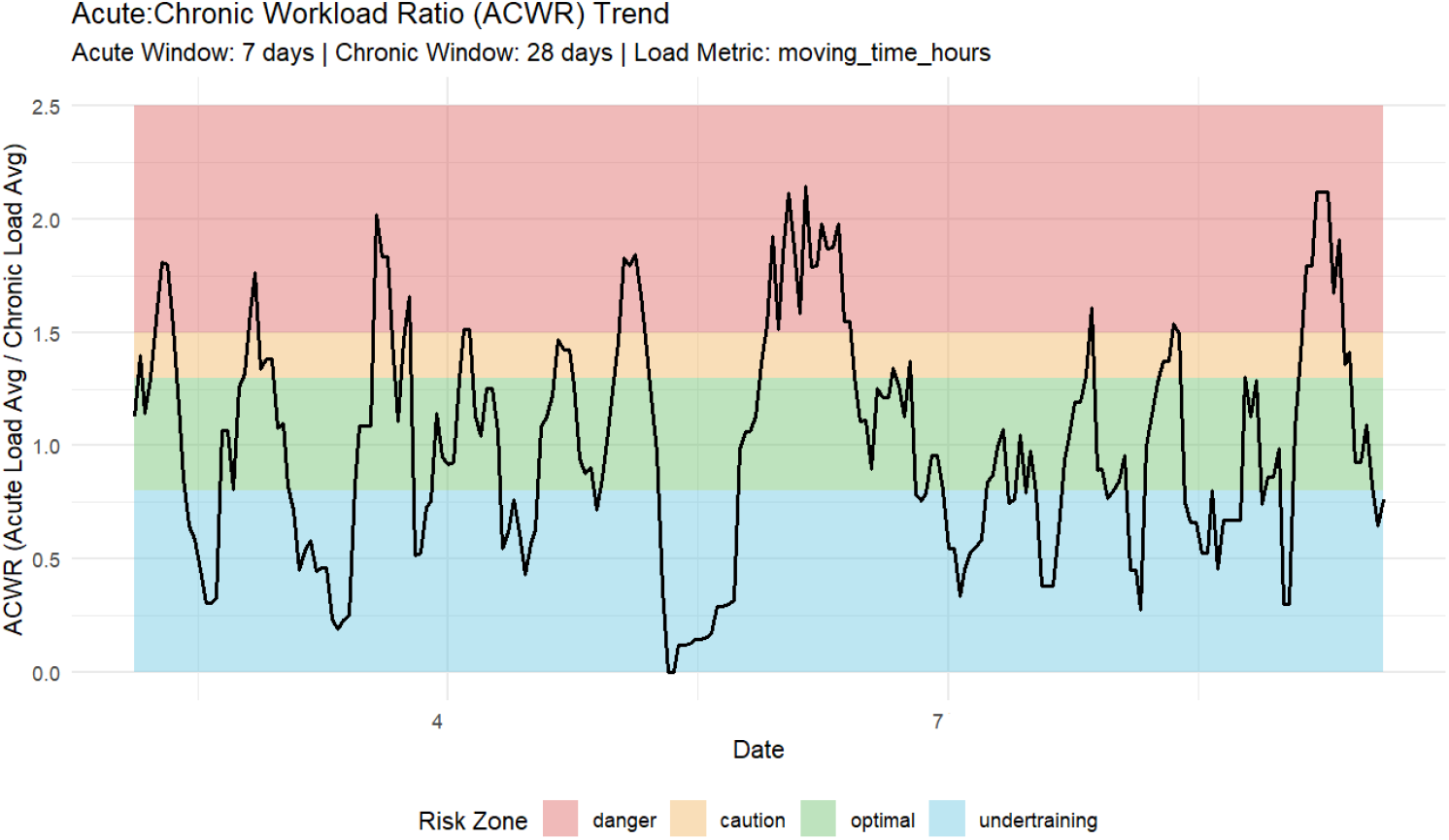
Example ACWR trend plot for running based on duration (Sample Data).

**Listing 3:**
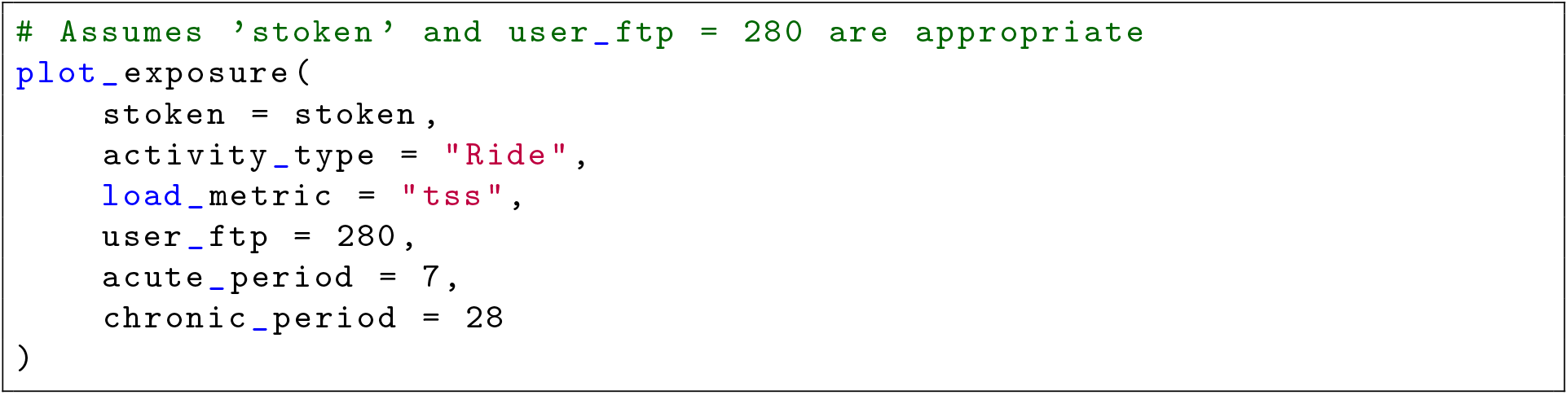
R code generating the Load Exposure plot (Figure 2).

**Figure 2:**
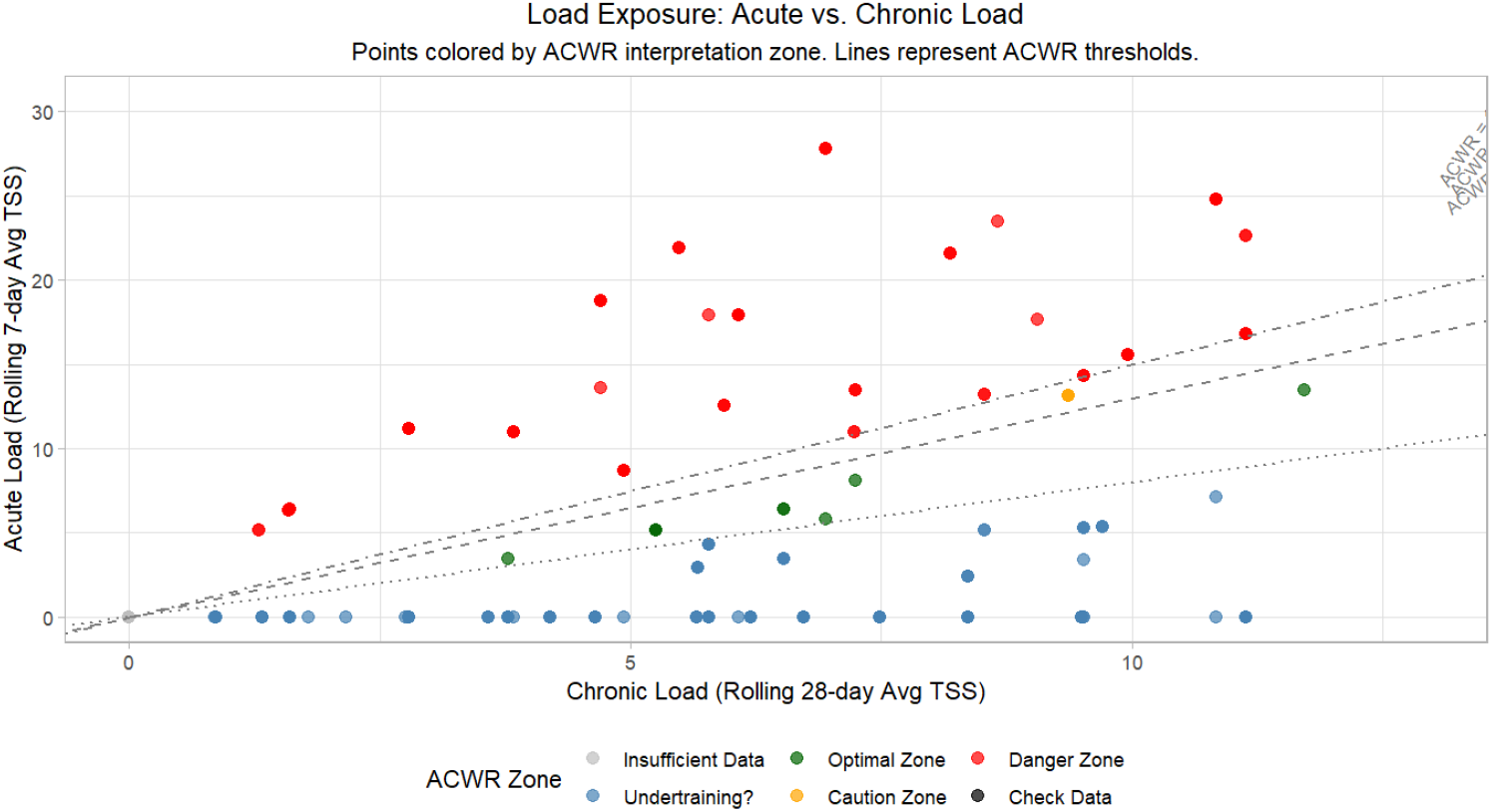
Example Load Exposure plot for cycling based on approximated TSS (FTP=280, Sample Data).

**Listing 4:**
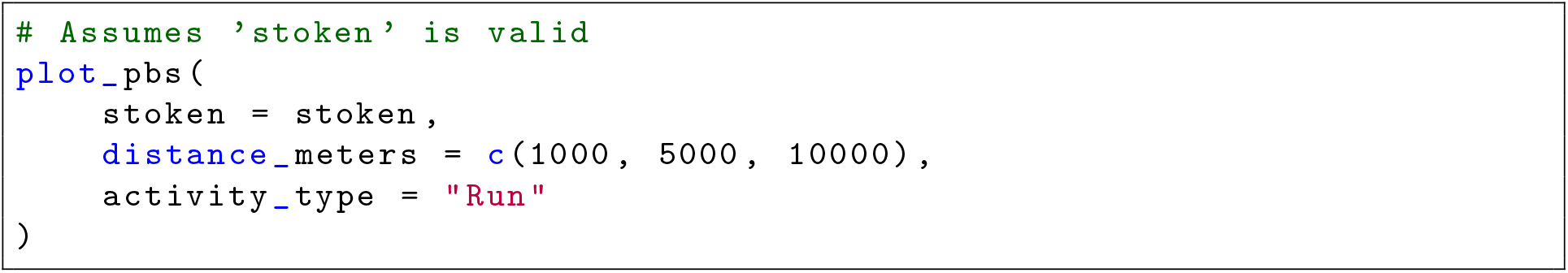
R code generating the Personal Bests progression plot (Figure 3).

**Figure 3:**
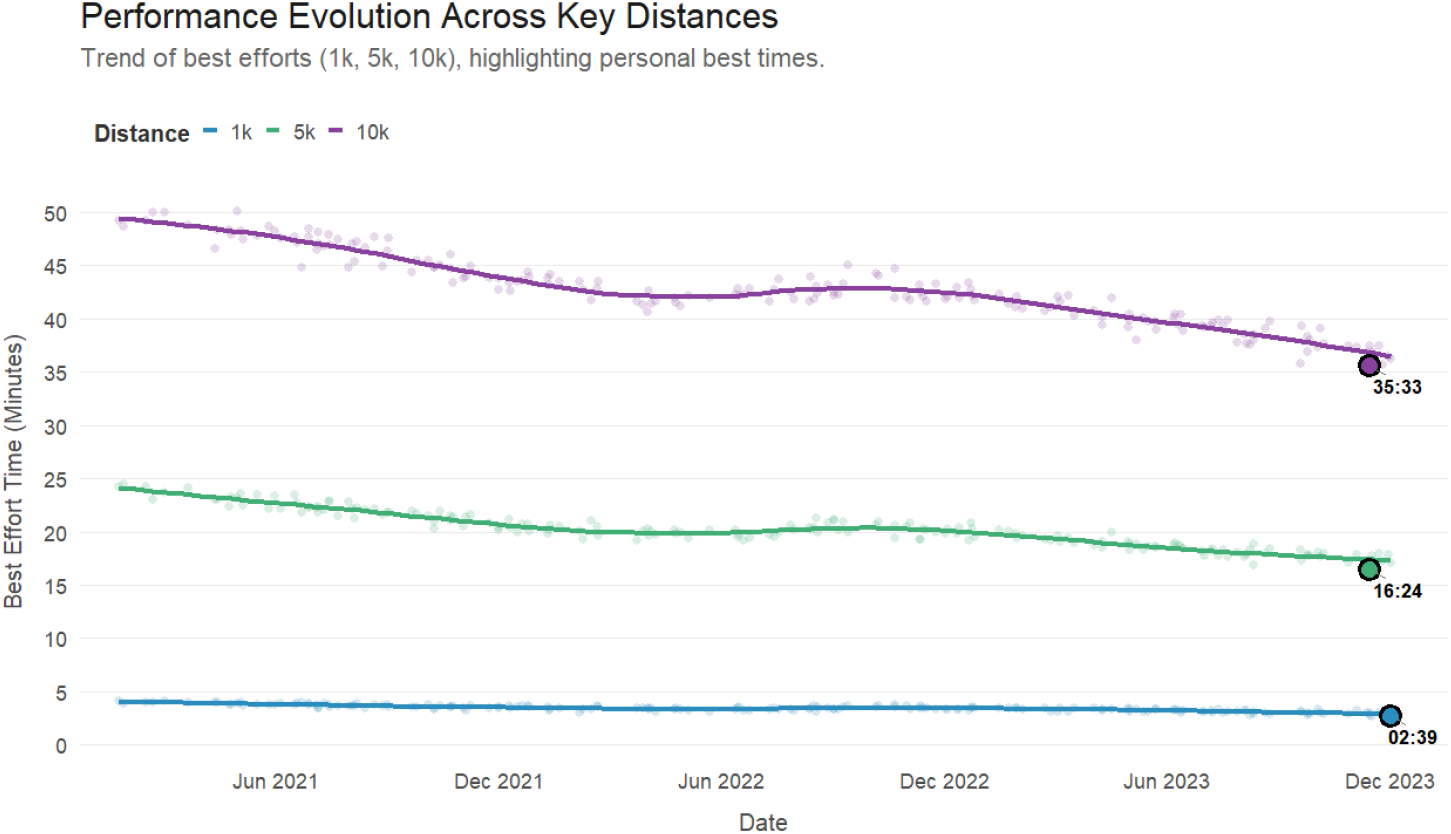
Example Personal Bests (PBs) progression based on Strava’s best_efforts data for 1km and 5km running (Sample Data).

**Listing 5:**
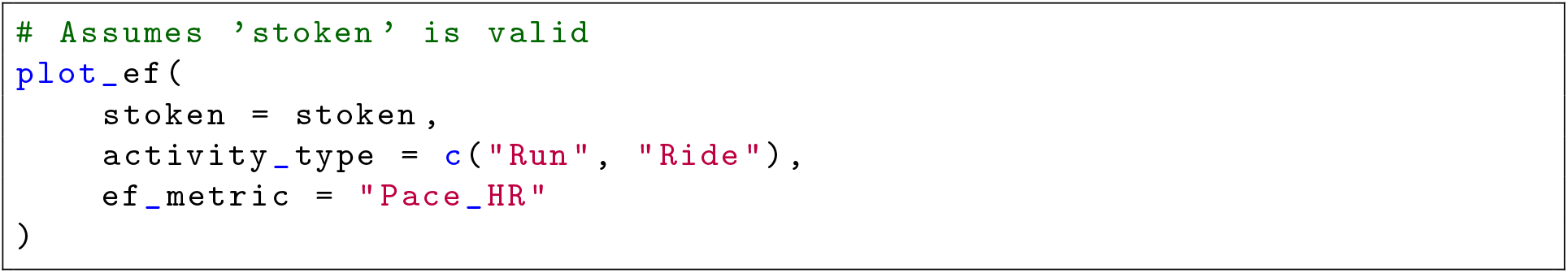
R code generating the Efficiency Factor trend plot (Figure 4).

**Figure 4:**
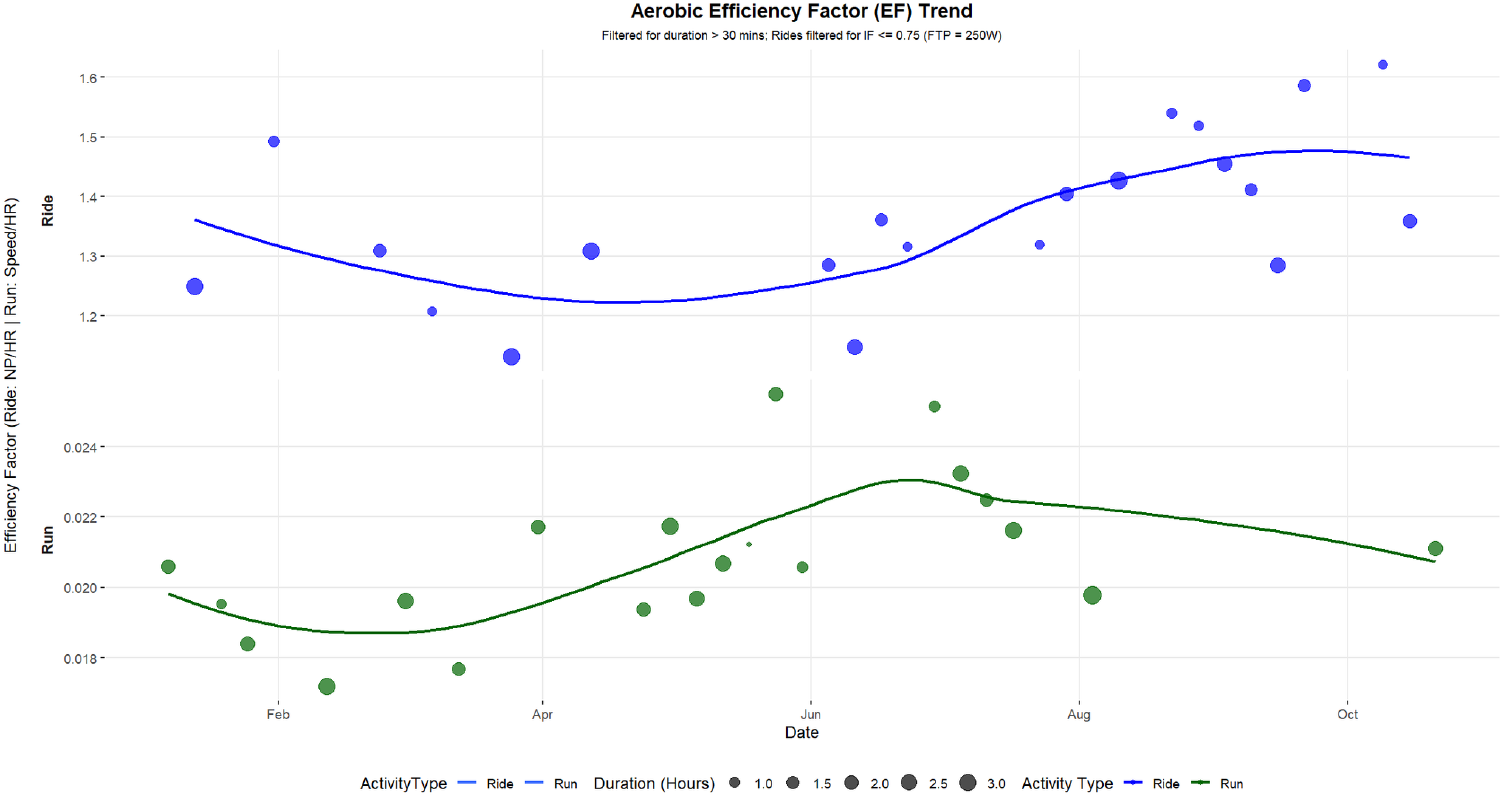
Example Efficiency Factor (EF) trend plot (Pace/HR) for running activities (>20 min, Sample Data).

**Listing 5:**
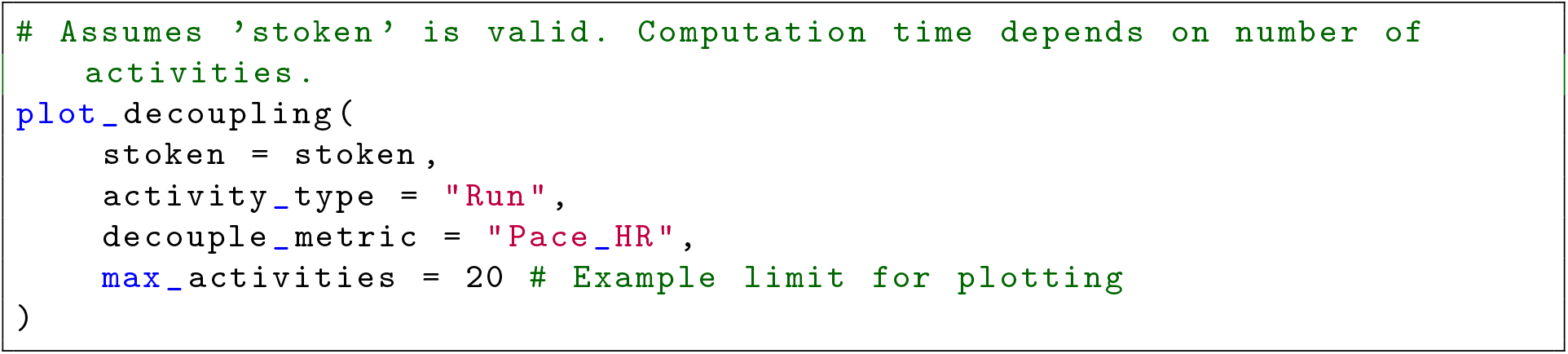
R code generating the Decoupling trend plot (Figure 5).

**Figure 5:**
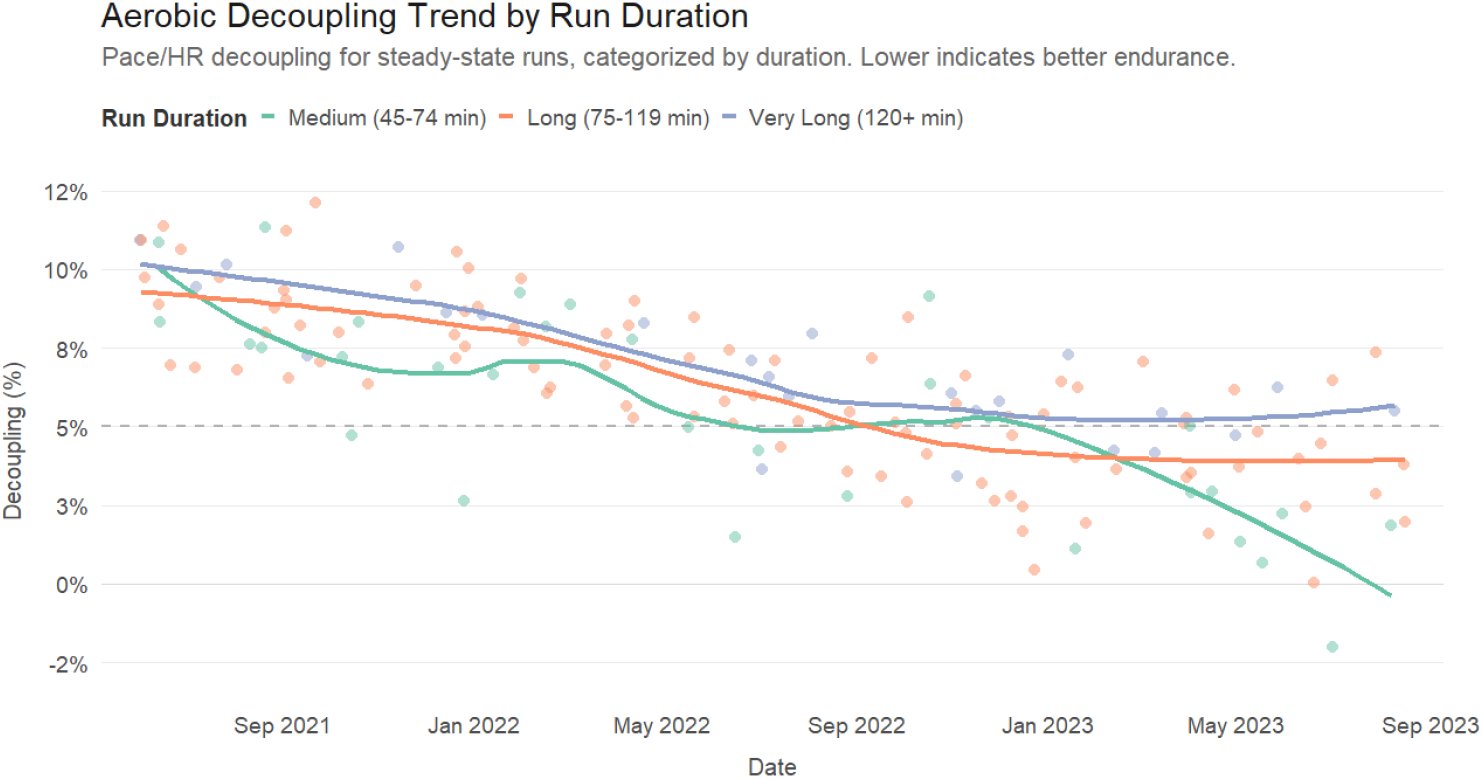
Example Decoupling trend plot (Pace/HR) for selected running activities (Sample Data).

## 4 Discussion

The `Athlytics` R package provides a specialized computational framework designed to address the practical challenges of analyzing key training load and physiological performance metrics derived directly from Strava API data. Its primary contribution lies in integrating data retrieval with standardized calculations and visualizations for established indicators (ACWR, Load Exposure, PBs, EF, Decoupling), offering a reproducible and efficient workflow within the R environment. This addresses a notable gap for researchers seeking to leverage ubiquitous wearable data sources without extensive bespoke programming.

### 4.1 Positioning within the Computational Methodology Landscape

`Athlytics` occupies a specific methodological niche relative to other R packages in the sports science and bioinformatics domains:

- Compared to `rStrava` [11], which provides essential data access, `Athlytics` adds a crucial analytical layer, transforming raw activity data into longitudinal physiological and performance metrics relevant to exercise science research questions.
- Unlike comprehensive file-based frameworks like `trackeR` [21], `Athlytics` is specifically optimized for the nuances and structure of API-sourced data (particularly from Strava) and provides pre-built modules for metrics often requiring custom implementation otherwise, facilitating a distinct and streamlined analytical workflow for this common data modality.
- Contrasting with single-metric packages like `ACWR` [22], `Athlytics` integrates ACWR analysis within a broader multi-metric framework, enabling the simultaneous investigation of load dynamics alongside indicators of physiological efficiency (EF) and fatigue (Decoupling), starting directly from data retrieval.
- By bundling these functionalities, `Athlytics` provides a unique, integrated solution that significantly lowers the barrier to entry for conducting specific types of longitudinal training analysis, thereby enhancing research feasibility and promoting methodological standardization.

### 4.2 Limitations and Implications for Quantitative Research

While `Athlytics` offers considerable convenience and workflow integration, its application in rigorous quantitative research necessitates a critical understanding of significant limitations arising from both the data source and the implemented calculation methods.

#### 1. Validity of Approximated Load Metrics (TSS/HRSS)

This is a fundamental limitation when using the `tss` or `hrss` options for `load_metric` in ACWR and Load Exposure calculations. `Athlytics` currently approximates these scores using activity summary statistics (average power/HR, duration, user-provided thresholds). This approach, while computationally pragmatic for API-derived summary data, fundamentally deviates from the theoretical basis of TSS (based on Normalized Power®) and HRSS (based on timein-zones), which are designed specifically to account for the non-linear physiological cost of intensity variations within a session [6, 10]. Using average values inherently masks the difference between, for instance, a steady-state aerobic session and an interval session with the same average power or heart rate but vastly different physiological impact. Consequently, ACWR or Load Exposure calculations based on these approximated scores may produce misleading results, particularly when comparing or aggregating activities with diverse intensity profiles. Such approximations are likely unsuitable for precise dose-response modeling, detailed physiological comparisons between training modalities, or research relying heavily on the accuracy of ACWR for injury prediction [3, 4]. Researchers employing these options must explicitly acknowledge this limitation and restrict interpretations primarily to very broad, potentially coarse-grained load trends, unsuitable for fine physiological inference. Future work aims to incorporate more accurate calculation methods (see Section 4.3).

#### 2. Reliability of Strava-Derived Data for Research

- **Personal Bests (PBs):** The PB tracking module relies entirely on Strava’s `best_efforts` field within activity summaries. The accuracy and completeness of this field are variable and depend on Strava’s internal processing and the quality of the underlying GPS data. GPS inaccuracies, variations in course measurement, and algorithmic artifacts mean these identified PBs may not represent true best performances over standardized distances. Using these data for rigorous performance tracking or inter-individual comparisons without external validation or careful data scrutiny is scientifically questionable.
- **General Data Quality:** Analyses are inherently dependent on the quality and completeness of the underlying Strava data accessed via the API. Missing sensor data (heart rate, power), GPS signal dropout or noise affecting pace and distance, and reliance on user-provided (and potentially inaccurate or outdated) FTP values for approximated TSS further compound the uncertainty. All results generated using `Athlytics` should therefore be interpreted conditionally upon the quality of the input data.

#### 3. Computational Scalability for Detailed Analyses

Obtaining detailed time-series data for decoupling analysis imposes higher computational costs and API quota usage, potentially limiting feasibility for studies involving very large datasets or frequent, nearreal-time monitoring.

#### 4. Constrained Scope of Physiological Inquiry

The current modules, while covering common metrics, omit more advanced physiological analyses (e.g., HRV, detailed metabolic modeling, sophisticated performance modeling like critical power). This focused scope provides valuable but incomplete insights into the complex physiological state of an athlete.

#### 5. Robustness and Error Handling

While functional, error handling for API issues or missing data fields could be further enhanced to provide more informative diagnostics for research workflows.

These limitations highlight the responsibility of the researcher to select appropriate analytical tools and metrics for their research context and to interpret results generated by `Athlytics` with a clear understanding of the underlying data and calculation methods.

### 4.3 Future Directions: Enhancing Research Capabilities

Future development efforts will focus on extending the framework’s capabilities and addressing limitations to better support quantitative exercise physiology research:

1. **Improving Quantitative Accuracy:** Prioritizing the implementation of optional workflows utilizing detailed stream data (when available and computationally acceptable) for more scientifically valid TSS/HRSS calculations based on established methods (e.g., Normalized Power®, time-in-zones) is crucial for enhancing analytical rigor for studies requiring higher fidelity load quantification.
2. **Expanding Methodological Repertoire:** Incorporating alternative established load metrics (e.g., TRIMP variants [10]) and performance models (e.g., basic critical power estimation from available data [23]) would enable comparative analyses and broader applicability.
3. **Refining Existing Analytical Modules:** Enhancing the robustness and providing options for verifying or manually curating PBs, and expanding visualization options for decoupling and other metrics.
4. **Facilitating Exploratory Analysis:** Investigating interactive components via frameworks like `shiny` [24] could enhance hypothesis generation and data exploration.
5. **Supporting Reproducible Science:** With `Athlytics` now available on CRAN, ongoing maintenance, continued enhancement of comprehensive documentation explicitly detailing limitations and assumptions, and adherence to software development best practices remain key priorities. Further development of vignettes that demonstrate robust research practices, including data quality assessment and appropriate metric selection/interpretation, will continue to enhance pedagogical and research value, ensuring the package remains a reliable tool for the scientific community.

## 5 Conclusion

The `Athlytics` R package provides a valuable **computational framework** designed to lower barriers and enhance the **reproducibility** of analyzing longitudinal training load and physiological metrics from ubiquitous wearable sensor data, specifically via the Strava API. By integrating data access with standardized calculations and visualizations for key indicators like ACWR, EF, and decoupling, it offers a streamlined and efficient workflow within the R environment, addressing a critical gap in the current analytical toolkit. While researchers must exercise significant caution regarding the inherent limitations, particularly the **scientific validity compromises associated with load metrics approximated from summary statistics** and the dependencies on Strava data quality, `Athlytics` represents a significant step towards facilitating more widespread, standardized, and efficient quantitative analysis in exercise physiology and sports science research leveraging real-world data. We anticipate this framework, especially with future enhancements addressing calculation fidelity, will serve as a useful tool for researchers and practitioners seeking initial data-driven insights into training responses, provided results are interpreted with appropriate scientific rigor.

## Acknowledgments

The development of the Athlytics package relied upon the R programming language ([9]) and benefited greatly from the contributions of the open-source R community through the various packages cited in this work. Access to the necessary data was made possible by the Application Programming Interface (API) provided by Strava, which is gratefully acknowledged.

